# Light microscopy of proteins in their ultrastructural context

**DOI:** 10.1101/2020.03.13.989756

**Authors:** Ons M’Saad, Joerg Bewersdorf

## Abstract

Resolving the distribution of specific proteins at the nanoscale in the structural context of the cell is a major challenge in fluorescence microscopy. Here we present a new concept that decrowds the intracellular space through 13 to 21-fold physical expansion while simultaneously retaining the proteins. This combination makes labeling of the proteome efficient enough that local protein densities are revealed and the cellular nanoarchitecture can be visualized by standard light microscopy.

## Main Text

Fluorescence microscopy has transformed the field of cell biology through its exceptional contrast and high specificity of labeling. With the advent of super-resolution microscopy, the three-dimensional (3D) distribution of proteins of interest can be imaged at spatial resolutions down to ~10 nm, revealing astounding sub-cellular organization at the nanoscale [1]. Showing these proteins in the ultrastructural context of the cell, however, has so far largely relied on correlative microscopy techniques that combine the resolving power and global contrast of electron microscopy (EM) with the information provided by high molecular specificity modalities, such as fluorescence microscopy. While correlative light/electron microscopy (CLEM) can yield information-rich images of the 3D landscape of the cell [2], it requires highly-specialized instruments and days to weeks of continuous data acquisition for a single 3D data set of a mammalian cell.

Here, we demonstrate a straightforward method of imaging whole-cell structural context based on Expansion Microscopy (ExM) [3–5]. In ExM, a biological sample is embedded and hybridized to a swellable polyacrylamide/sodium polyacrylate copolymer. By absorbing water, the gel physically expands by a factor of ~4 in all three dimensions. In iterative expansion microscopy (iExM), iteratively anchoring fluorescent labels twice to hydrogel networks has yielded expansion factors of typically up to 20 [6]. The commonly used proteases, which digest proteins to allow for homogeneous expansion, however, result in the degradation of most cellular content. Variants of ExM such as Magnified Analysis of the Proteome (MAP) and Ultrastructure-ExM (U-ExM) have addressed this problem by preventing inter- and intra-protein crosslinks during the tissue-hydrogel hybridization step and by using anionic surfactants and heat to isotropically separate neighboring proteins [7, 8]. However, despite their promise, these techniques have been limited to an expansion factor of ~4 as they are not compatible with iterative expansion.

We hypothesized that by embedding a dense expanded sample prepared with a cleavable crosslinker in a second dense superabsorbent hydrogel, entanglements between polymer chains of the first and final hydrogels will physically interlock protein-polymer hybrids in this latter polymer network, thereby preserving the proteome while iteratively expanding it. This type of polymer entanglement, first described by Sam Edwards in 1967 [9], would retain most of the cellular proteome in the final hydrogel, while simultaneously expanding the sample by about a factor of 4^2^. The hydrogel chemistry employed in this method is reminiscent of semi-interpenetrating polymer networks (semi-IPNs), where one polymer is crosslinked and the second is not [10]. This method combined with global fluorescent labeling, which targets all separated proteins, would reveal the overall landscape of the cell with a light microscope - resembling the contrast of heavy-metal EM stains.

Based on this concept, we developed an ExM method which we named, in reference to the philosophy of labeling the whole (Greek: pan) proteome, pan-ExM. In brief (**Supplementary Fig. 1**), previously fixed cells cultured on coverslips are incubated in a solution of acrylamide (AAm) and formaldehyde (FA) to simultaneously prevent inter-protein crosslinks and tether proteins to the hydrogel. Next, the cells are embedded in a polyacrylamide/polyacrylate co-polymer cross-linked with N,N’-(1,2-dihydroxyethylene)bisacrylamide (DHEBA), an acrylamide crosslinker with a cleavable diol bond. After polymerization, the sample is delipidated and denatured with sodium dodecyl sulfate (SDS) and heat. The hydrogel is then expanded ~4-fold in deionized water and embedded in a neutral polyacrylamide hydrogel cross-linked with DHEBA to maintain the gel in its expanded state during the subsequent treatments. Afterwards, the gel is incubated in a solution of FA and AAm for additional anchoring of previously masked primary amines to the next hydrogel. This composite hydrogel is embedded in a third hydrogel, a polyacrylamide/polyacrylate co-polymer cross-linked with N,N-methylenebis(acrylamide) (BIS), a non-cleavable acrylamide crosslinker. Next, sodium hydroxide is used to cleave the crosslinks of the first and second hydrogels and the final hydrogel is expanded in deionized water a second time resulting in a linear expansion factor of ~13-21 (depending on the gel chemistry) as measured by the size of HeLa cell nuclei before and after expansion (**Supplementary Table**).

Typical organic dyes are ~1 nm in diameter and thus comparable in size to the distance between proteins in the densely crowded cellular interior [11]. We rationalized that the expansion of the sample will lead to substantial molecular decrowding and will reveal previously masked binding sites and avoid steric hindrance of fluorescent labels. To test this concept, we labeled cells with NHS ester-dye conjugates, taking advantage of the abundance of primary amines on proteins. A visual comparison of HeLa cells non-expanded, expanded once, or twice, imaged with a standard confocal microscope (**Fig. 1a-c; Supplementary Fig. 2**), confirms the validity of this concept: non-expanded cells show essentially a uniform staining, revealing little information. Single-expanded cells start to show the outlines of organelles such as mitochondria, but expanding cells twice by an overall factor of ~13 to 21, in contrast, allows to spatially separate organelles which originally were less than 20 nm apart by a standard diffraction-limited (~250-nm resolution) confocal microscope. Analogous to EM, now resolvable hallmark features such as mitochondrial cristae (**Fig. 1d**) or the stacking of Golgi cisternae (**Fig. 1c inset**) allow for the identification of organelles by their morphological characteristics.

**Fig. 1:**
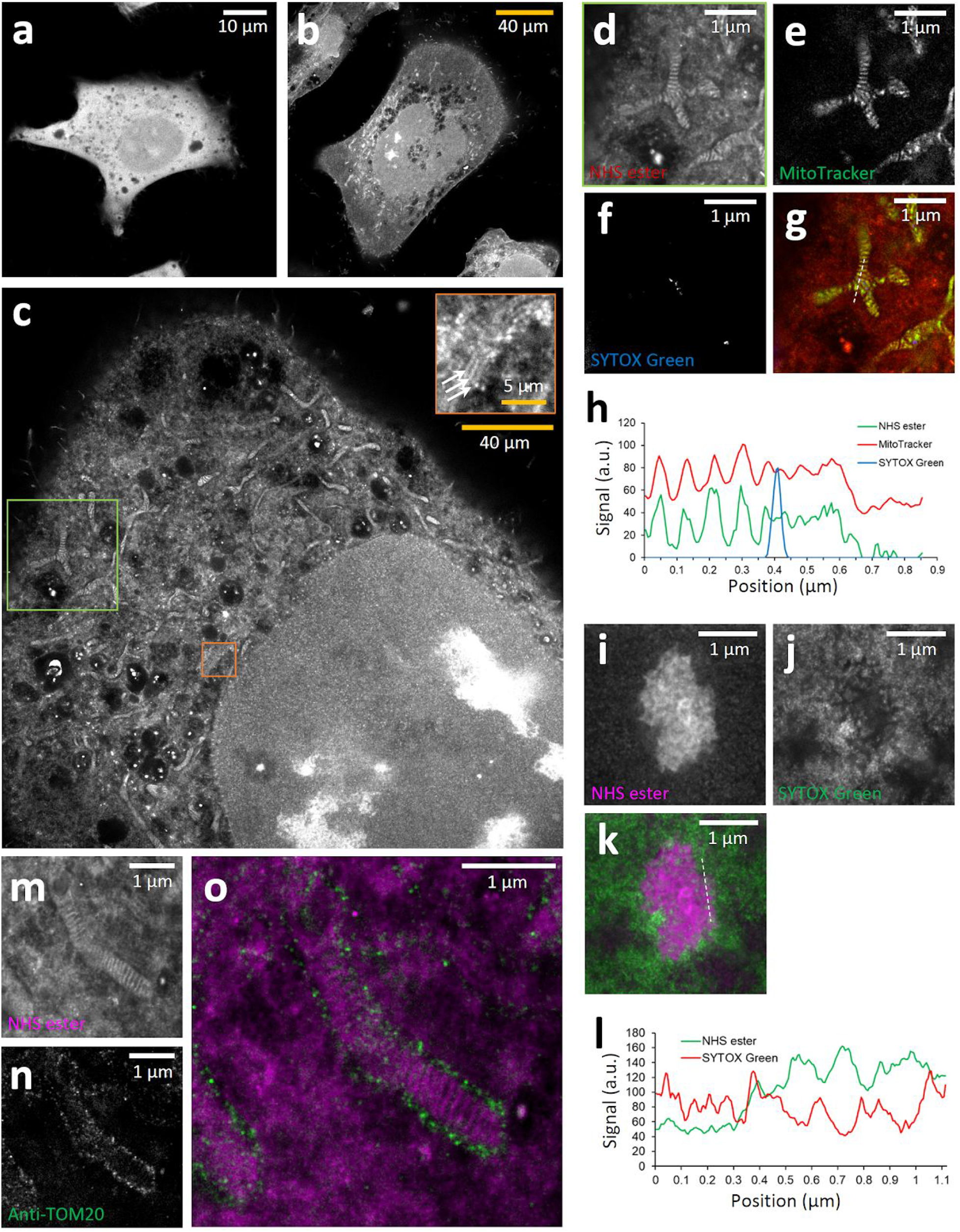
pan-ExM is compatible with standard fluorescent labeling techniques. **a**, Non-expanded HeLa cell. **b**, HeLa cell expanded once. **c**, pan-ExM expanded HeLa cell. The area in the orange box is shown in inset, revealing Golgi cisternae (arrow heads). The cells in **a-c** were labeled with NHS ester, **d**, magnified green box in **c**, showing cristae in mitochondria. **e**, MitoTracker Orange stain in the same area. **f**, SYTOX Green stain showing DNA in mitochondrial nucleoids. **g**, overlay of **d-f**. **h**, cross-section along the dashed line shown in **g**. **i**, NHS-ester rich area in a HeLa cell nucleus showing a nucleolus. j, SYTOX Green DNA stain corresponding to area shown in **i**. **k**, overlay of **i-j**. **l**, cross-section along the dashed line shown in **k**. **m**, NHS ester-labeled mitochondrion. **n**, same area as in **m**, showing anti-TOM20 immunostaining and revealing the outer membrane of the mitochondrion. **o**, overlay of **m-n**. White scale bars show expansion-corrected values. Yellow scale bars are not corrected for the expansion factor.

Importantly, our protocol is compatible with well-established staining methods such as MitoTracker Orange, a live-cell dye which localizes to mitochondria (**Fig. 1e,g**) and DNA-intercalating dyes such as SYTOX Green (**Fig. 1f,j**). Moreover, antibody-labeling of the outer mitochondrial membrane protein TOM20 (**Fig. 1n,o**) demonstrates the compatibility of pan-ExM with immunohistochemistry. A comparison of fluorescence signal levels of antibodies applied without expansion and after one or two expansion steps shows no decrease in signal (**Supplementary Figure 3**). This confirms the good protein retention capabilities of our protocol.

To investigate structural preservation of features at the sub-organelle scale, we quantified the distance between mitochondrial cristae (**Fig. 2a,b**). The determined expansion-corrected distance of 85 ± 22 nm (mean ± s.d.) is in good agreement with previously published observations in living HeLa cells [12]. While the endoplasmic reticulum (ER) did not show a characteristic NHS-ester staining that would make it directly identifiable, we could visualize it by overexpressing ER-membrane localized Sec61β-GFP and immunolabeling it with an anti-GFP antibody (**Fig. 2c-f**). The expansion-corrected diameter of these tubules (47 ± 10 nm, mean ± s.d.; **Fig. 2g**) was slightly smaller than diameters determined by super-resolution light microscopy [13] but consistent with diameters of ER tubules determined by EM [14]. We overexpressed the Golgi protein mannosidase II (ManII) fused to GFP, labeled it with an anti-GFP antibody after applying our pan-ExM protocol, and recorded a 3D image stack with a confocal microscope. The immunolabeled Golgi cisternae overlayed well with sheet-like structures visible in the NHS-ester image (**Fig. 2h-s, Supplementary Video 1**). Imaging at different axial positions revealed the intricate structure of the convoluted Golgi ribbon.

**Fig. 2:**
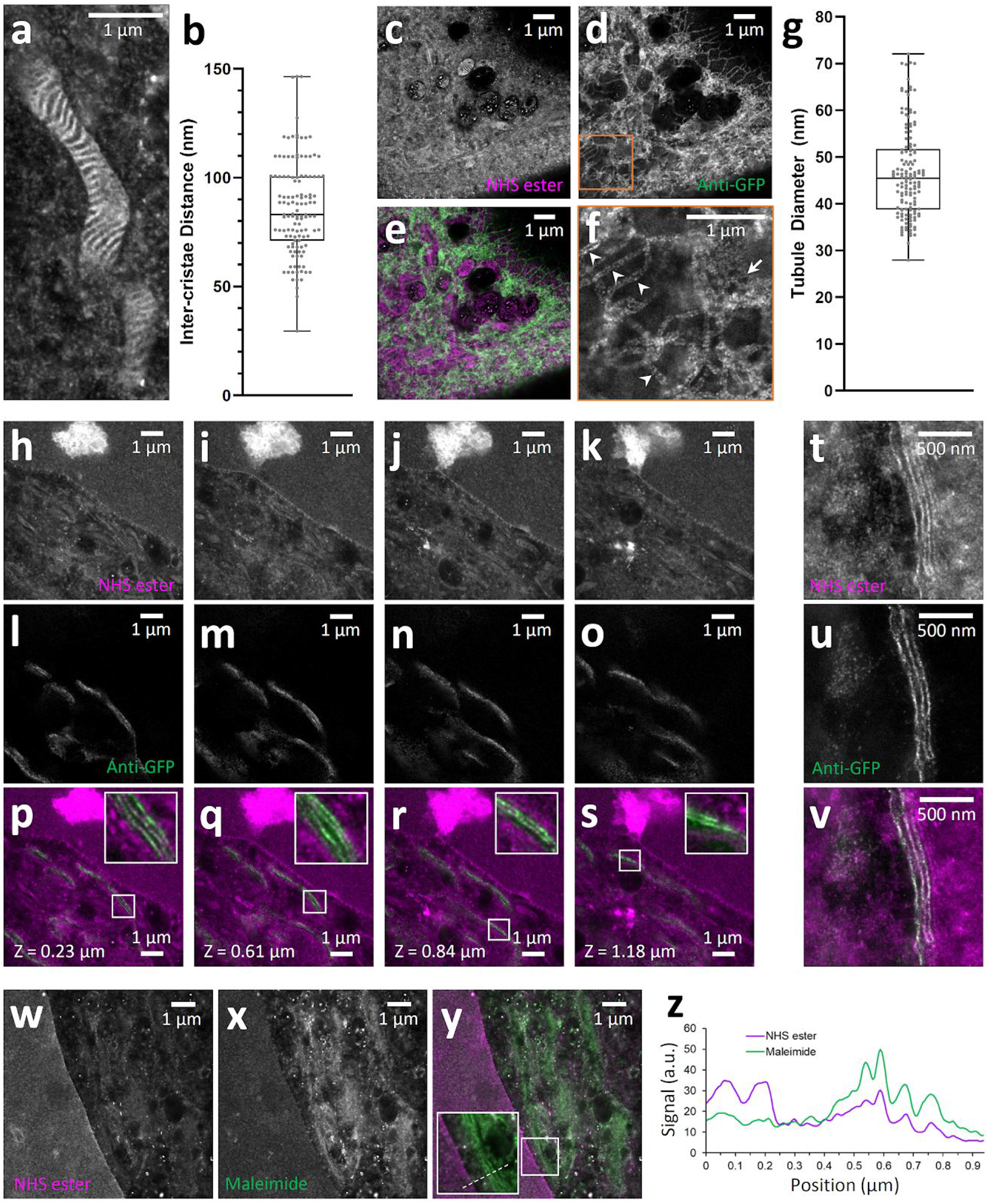
pan-ExM reveals ultrastructural details of cellular organelles. **a**, NHS-ester labeled mitochondrion in a HeLa cell revealing cristae. **b**, distance distribution between neighboring cristae (n = 123, N = 4 experiments). **c**, NHS ester channel of a HeLa cell expressing ER-membrane localized Sec61β-GFP. **d**, anti-GFP label in the same area revealing the ER. **e**,overlay of **c-d**. **f**, zoomed-in anti-GFP image of the area denoted by the orange box in **d**, revealing individual ER tubules, clearly resolved as hollow tubules (arrow heads), and a dense network of ER tubules (arrow). **g**, distribution of ER tubule diameters (n = 142, N = 2 cells). **h-s**, images from a 3D image stack featuring the Golgi complex next to the nucleus of a HeLa cell expressing Golgi-localized ManII-GFP. **h-k**, NHS ester images at axial positions 0.23, 0.61, 0.84 and 1.18 μm, respectively. **l-o**, anti-GFP images of the same fields of view as **h-k. p-s**, overlay of **h-o**. **t**, STED super-resolution image showing the NHS ester channel of a Golgi stack in a ManII-GFP expressing HeLa cell. **u**, anti-GFP STED image of the same area. **v**, overlay of **t-u**. **w-y**, nuclear periphery of a HeLa cell (nucleus on the left) labeled with NHS ester and maleimide, revealing the cysteine-enriched Golgi complex. **w**, NHS ester channel. **x**, maleimide channel. **y**, overlay of **w-x**. The inset shows the zoomed-in white box. **z**, line profile along the dashed line in **y**, revealing the change in NHS ester to maleimide staining across the Golgi and nucleus. All scales are corrected for the determined expansion factor. For box plots, median and interquartile range are shown with whiskers drawn down to the minimum and maximum values.

To allow a more detailed view of the distribution of ManII in the Golgi, we imaged expanded samples with a STED super-resolution microscope. It is known that ManII is located primarily in the medial cisternae of the Golgi and, when overexpressed, also in cis cisternae [1]. Consistent with this observation, we find the ManII stain to highlight 2 to 3 cisternae located at one side of the Golgi stack (**Fig. 2t-v**).

The data presented here demonstrates, from our perspective, pan-ExM’s potential to revolutionize light microscopic imaging of cellular structures. Pan-ExM combines, (i), 13 to 21-fold linear expansion (obtained through iterative expansion), (ii), the protein retention capabilities of MAP and U-ExM, and (iii) the transfer of entrapped proteins to the final expansion hydrogel by polymer chain entanglement. While these individual techniques have demonstrated merit, we see their true power in their combination with labeling the whole proteome: staining the cell with NHS esters shifts from a largely pointless exercise in a non-expanded cell (**Fig. 1a,b**) to a technique with EM-like contrast, capable of revealing nanoscopic structural hallmarks that allow users to identify organelles without the need for specific staining. While the obtained resolution does not reach the level of EM, its straightforward combination with single, or multiple, specific labels stands in stark contrast to the complex protocols of immuno-gold EM or CLEM. Our sample preparation is completed in a few days; subsequent imaging takes less than a minute (2D image) to tens of minutes (3D data set; **Supplementary Video 2**) per cell on a conventional light microscope. We anticipate that light-sheet microscopes or other instruments optimized for ExM [15] will accelerate future pan-ExM data acquisition substantially.

We have shown here using anti-GFP and anti-TOM20 antibodies that pan-ExM is compatible with immunolabeling. However, many of the antibodies we tested did not work. We anticipate that this caveat can be overcome in the future by (i) focusing on antibodies which have been shown to work well in western blots due to the commonality of the denaturing step and (ii) avoiding the high basic pH currently used to cleave the DHEBA-formed crosslinks before the final expansion by switching to a chemical crosslinker that can be dissolved at more physiological pH values such as N,N’-diallyl-tartardiamide (DATD).

It remains to be investigated down to which size scale structures are preserved in pan-ExM. Based on our protocol, we expect that proteins lose their tertiary structure and unfold during the sample preparation process. At the tens-of-nanometer scale, our data shows that membrane-bound organelles such as mitochondria, the Golgi complex and the ER are represented well after expansion. In the other extreme: lipid droplets, having a lipid core, are dissolved by the used detergents, leaving only a shadow image of their former shape. Future systematic studies, refinements to the protocols and engineered probes, such as cross-linkable lipid labels [16] and reversible protein crosslinking reagents, will answer these questions and promise further improvements.

On the positive side, pan-ExM leads to substantial molecular decrowding of the sample: the average distance of 3 nm between proteins in the native cell becomes about 50 nm after expansion and the space in between proteins becomes easily accessible by probes even as large as antibodies (6-12 nm in size). Furthermore, the expansion protocol most likely disrupts protein clusters through denaturation, revealing previously inaccessible binding sites. It is worth pointing out that, in contrast to optical super-resolution microscopy techniques which are ultimately limited in resolution by the size of the fluorescent labels, ExM is not constrained by label-size (if labeling is applied after expansion). Moreover, problems with labeling density which are a major limitation in optical super-resolution microscopy get alleviated: steric hindrance and fluorescence quenching in densely labeled samples become essentially irrelevant after 13-to 21-fold expansion. On the contrary, the created gaps between endogenous molecules provide space for potential biochemical amplification of the label, boosting the sensitivity of the technology.

The fact that many organelles can easily be identified by eye through their characteristic NHS-ester stain suggests that pan-ExM is well-suited for automated segmentation and classification algorithms to identify organelles of interest. Training of machine learning algorithms can be facilitated by additional imaging channels showing specific stains of the target organelle, for example MitoTracker for mitochondria, during the learning phase.

Pan-staining is not restricted to NHS ester: dye-conjugated maleimide, for example, can reveal cellular domains with high cysteine-rich protein content, as shown in **Fig. 2x** for the Golgi complex, most likely related to the high levels of palmitoyltransferases in the Golgi which have cysteine-rich domains [17]. The combination with NHS ester labeling (**Fig. 2w**) shows intriguing patterns of differential staining (**Fig. 2y,z**) which provide information well beyond the monochromatic, EM-like contrast achievable with a single pan-staining - analogous to Haemotoxylin and Eosin (H&E) staining but on the nanoscale. This approach can be easily generalized to other sub-proteomes like phosphorylated, glycosylated or palmitoylated (**Supplementary Fig. 4**) proteins. We believe that the combination of two or more of these pan-stainings has great potential for automated segmentation of organelles and other subcellular structures of interest, and can also help to answer how different sub-proteomes are distributed in the cell. We emphasize that the success of our pan-stain approach fundamentally benefits from two key strengths: high levels of protein retention and molecular decrowding provide access to large numbers of target sites; high levels of expansion offer the effective spatial resolution required to identify structures and their cellular context.

## Acknowledgements

We would like to thank Dr. James Rothman and Dr. Andreas Ernst for fruitful discussions. We are grateful to Lena Schroeder for her guidance on fluorescent staining methods and to Zach Marin and Dr. Kenny Chung for comments on the manuscript. This work was supported by grants from the Wellcome Trust (203285/B/16/Z) and NIH (P30 DK045735, S10 OD020142).

## Author Contributions

O.M. developed the concept and sample preparation protocols and prepared the samples. O.M. and J.B. performed the imaging experiments and analyzed the data. J.B. provided guidance on the project. O.M. and J.B. wrote the manuscript.

## Competing Interests

J.B. has financial interests in Bruker Corp. and Hamamatsu Photonics. The authors filed a patent application covering the presented method.

## Data availability

The datasets generated and/or analyzed during the current study are available from the corresponding author on reasonable request.

## Methods

### Coverglass preparation

Before plating HeLa cells, 12-mm round glass coverslips (Electron Microscopy Sciences, catalog no. 72230-01) were cleaned in a sonic bath (Bronson) submerged in 1M KOH (Macron Fine Chemicals; catalog no. 6984-04) for 15 minutes and then rinsed with MilliQ water three times. Glass was then sterilized with 100% ethanol, incubated with 5 μg/mL fibronectin (Sigma-Aldrich, catalog no. F1141) for 30 minutes, and rinsed with sterile phosphate buffered saline (PBS; Gibco, catalog no. 10010023) before adding media and cells.

### Cell culture

HeLa cells were grown in Dulbecco’s modified Eagle medium (DMEM; Gibco, catalog no. 21063029), supplemented with 10% fetal bovine serum (FBS; Gibco, catalog no. 10438026), and 1% mL/L penicillin-streptomycin (Gibco, catalog no. 15140122) at 37 °C with 5% CO_2_. Cells were passaged twice to three times a week and used between passage number 2 and 20. Passaging was performed using 1× PBS and 0.05% Trypsin-EDTA (Gibco, catalog no. 25300054). Approximately 24 hours before fixation, cells were seeded on fibronectin-coated glass coverslips at 65,000 cells per well.

### Plasmids

For labeling the medial Golgi, GFP-ManII was expressed. GFP-ManII was made from pEGFP-N1 (Takara Bio Inc.) to include amino acids 1-137 of mouse MAN2A1 fused to GFP, such that GFP is expressed in the Golgi lumen. For labeling the ER membrane, mEmerald-Sec61-C-18 was expressed. mEmerald-Sec61-C-18 was a gift from Michael Davidson (deceased, formerly Florida State University, Tallahassee, FL; Addgene plasmid # 54249; referred to as GFP-Sec61β).

### Transfection

GFP-ManII and GFP-Sec61β expression in HeLa cells used DNA transfection by electroporation as detailed in [1]. Briefly, DNA was introduced into the cells using a NEPA GENE electroporation device. Approximately 1 million cells were rinsed in Opti-MEM (Gibco, catalog no. 31985070) and then resuspended in Opti-MEM with 10 μg DNA in an electroporation cuvette with a 2-mm gap (Bulldog Bio, catalog no. 12358346). Cells were electroporated with a poring pulse of 125 V, 3-ms pulse length, 50-ms pulse interval, 2 pulses, with decay rate of 10% and + polarity; followed by a transfer pulse of 25 V, 50-ms pulse length, 50-ms pulse interval, 5 pulses, with a decay rate of 40% and ± polarity. After electroporation, the cells were seeded on fibronectin-coated coverslips (see coverglass preparation section). Samples were fixed 24 to 36 hours after electroporation.

### MitoTracker Orange staining

Live cells were incubated with 0.5 μM of MitoTracker Orange CMTMRos (Invitrogen, catalog no. M7510) for 30 min at 37°C and 0.5% CO_2_. Next, the cells were washed three times with cell media and fixed immediately after (see cell fixation section).

### Cell fixation

Cells were fixed with 3% formaldehyde (FA) and 0.1% glutaraldehyde (GA) (Electron Microscopy Sciences, catalog nos. 15710 and 16019 respectively) in 1× PBS for 15 minutes at RT. Afterwards, samples were rinsed three times with 1× PBS.

### Pan-ExM protocol

#### Reagents

Acrylamide (AAm; catalog no. A9099), N,N’-(1,2-dihydroxyethylene)bisacrylamide (DHEBA; catalog no. 294381), N,N’-Cystaminebisacrylamide (BAC; catalog no. 9809) were purchased from Sigma-Aldrich. Three different batches of sodium acrylate (SA) were used. The first batch (catalog no. 408220, lot no. MKCF0390) was purchased from Sigma-Aldrich. The second and third batches (catalog no. sc-236893C, lot nos. H3019 and L0619) were purchased from Santa Cruz Biotechnology. N,N-methylenebis(acrylamide) (BIS; catalog no. J66710) was purchased from Alfa Aesar. Ammonium persulfate (APS; catalog no. AB00112), N,N,N’,N’-tetramethylethylenediamine (TEMED; catalog no. AB02020), tris [hyroxymethyl] aminomethane (Tris; catalog no. AB02000), and 20% sodium dodecyl sulfate solution in water (SDS; AB01922) were purchased from American Bio. Sodium chloride (NaCl; catalog no. 3624-01) was purchased from J.T. Baker.

#### Gelation chamber

The gelation chamber was constructed using a glass microscope slide (Sigma-Aldrich, catalog no. S8400) and two spacers each consisting of a stack of two N1.5 22 x 22 mm coverglass (Fisher Scientific, catalog no. 12-541B) glued on both sides of the cell-adhered coverslip. N1.5 22 x 22 mm coverglass was used as a lid after adding the monomer solution. This geometry yielded an initial gel thickness size of ~170 μm.

#### First-round of expansion

HeLa cells, previously fixed as described in the “Cell fixation”section, were incubated in post-fix solution (0.7% FA + 1% AAm in 1× PBS) for 6 to 7 hours at 37°C. Next, the cells were washed twice with 1× PBS and embedded in first hydrogel monomer solution (19% (w/w) SA + 10% AAm (w/w) + 0.1% (w/w) DHEBA + 0.25% (v/w) TEMED + 0.25% (w/w) APS in 1× PBS). Gelation proceeded first for 15 min at room temperature (RT) and then 1.5 hours at 37°C in a humidified chamber. Coverslips with hydrogels were then incubated in ~1 mL denaturation buffer (200 mM SDS + 200 mM NaCl + 50 mM Tris in MilliQ water, pH 6.8) in 35 mm dishes for 15 min at RT. Gels were then transferred into denaturation buffer-filled 1.5 mL Eppendorf tubes and incubated at 73 °C for 1 hour. Next, gels were placed in petri dishes filled with MilliQ water for the first expansion. Water was exchanged at least twice every 1 hour and then the gels were incubated overnight in MilliQ water. Gels expanded between 4.0× and 4.5× according to SA purity.

#### Re-embedding in neutral hydrogel

Expanded hydrogels were incubated in a fresh re-embedding monomer solution (10% (w/w) AAm + 0.05% (w/w) DHEBA + 0.05% (v/w) TEMED + 0.05 (w/w) APS in 1× PBS) three times for 20 min on a rocking platform at RT. Immediately after, gels were placed between two pieces of no. 1.5 coverglass and incubated at 37 °C for 1.5 hours in a nitrogen-filled chamber. Next, gels were detached from the coverglass and washed three times with 1× PBS for 30 minutes each at RT. Gels were incubated in post-fix solution (0.7% FA + 1% AAm in 1× PBS) for 15 min at RT and then for 9 hours at 37°C.

#### Second-round of expansion

Re-embedded hydrogels were incubated in a fresh second hydrogel monomer solution (19% (w/w) SA + 10% AAm (w/w) + 0.1% (w/w) BIS + 0.05% (v/w) TEMED + 0.05% (w/w) APS in 1× PBS) four times for 15 min each on a rocking platform on ice. Immediately after, gels were placed between two pieces of no. 1.5 coverglass and incubated at 37 °C for 2 hours in a nitrogen-filled chamber. To dissolve DHEBA crosslinks, gels were incubated in 0.2 M NaOH for 1 hour on a rocker. Gels were next detached from the coverglass and washed three times with 1× PBS for 30 minutes each. Subsequently, the gels were labeled with antibodies and stained with NHS ester dyes. Finally, the gels were placed in petri dishes filled with MilliQ water for the second expansion. Water was exchanged at least twice every 1 hour at RT, and then the gels were incubated overnight in MilliQ water. Gels expanded between 4.0× and 4.5× according to SA purity for a final expansion factor of 15× to 20×. For the ER samples, the cleavable crosslinker BAC was used instead of BIS at a concentration of 0.1% (w/w).

### Antibody labeling after pan-ExM

For both Golgi and ER samples, gels were incubated for 36 to 40 hours at RT on a rocking platform with polyclonal rabbit anti-GFP antibody (Invitrogen, catalog no. A11122) diluted to 1:250 in antibody dilution buffer (2% BSA in 1× PBS). For mitochondria samples, gels were incubated 24 hours at RT on a rocking platform with rabbit anti-TOM20 antibody (Abcam, catalog no. ab78547). Gels were then washed in PBS-T (0.1% Tween 20 in 1× PBS) three times for 20 minutes each. Next, the samples were incubated for 24 hours at RT on a rocking platform with ATTO647N-conjugated anti-rabbit antibodies (Sigma-Aldrich, catalog no. 40839) diluted to 1:250 in the antibody dilution buffer. The gels were subsequently washed in PBS-T three times for 20 minutes each. Finally, the samples were rinsed and stored in 1× PBS at RT until subsequent treatments. Note that bovine serum albumin (BSA; catalog no. 001-000-162) was purchased from Jackson ImmunoResearch and Tween 20 (catalog no. P7949) was ordered from Sigma-Aldrich.

### NHS ester pan-staining after pan-ExM

After antibody labeling, gels were incubated for 1.5 hours at RT on a rocking platform with either 20 μg/mL NHS ester-ATTO594 (Sigma-Aldrich, catalog no. 08741) or 20 μg/mL NHS ester-ATTO532 (Sigma-Aldrich, catalog no. 88793) or 200 μM NHS ester-DY634 (Dyomics, catalog no. 634-01A), dissolved in 100 mM sodium bicarbonate (Sigma-Aldrich, catalog no. SLBX3650) buffer. The gels were subsequently washed five times in 1× PBS for 20 minutes each. Note that for the experiment where HeLa cells expanded never, once, or twice with pan-ExM were compared visually, the same concentration of NHS ester-ATTO594 and labeling conditions were used.

### Maleimide pan-staining after pan-ExM

Gels were reduced with 50 mM Tris(2-carboxyethyl)phosphine hydrochloride solution (TCEP) (Sigma-Aldrich, catalog no. 646547) in 1× PBS for 30 minutes at RT and subsequently incubated for 1.5 hours in an inert environment with 20 μg/mL maleimide-ATTO594 (Sigma-Aldrich, catalog no. 08717) dissolved in deoxygenated 150 mM Tris-Cl pH 7.4 solution. The gels were then washed three times in 1× PBS for 20 minutes each.

### Sytox Green staining after pan-ExM

Gels were incubated with Sytox Green (Invitrogen, catalog no. S7020) diluted 1:3,000 in calcium- and magnesium-free HBSS buffer (Gibco, catalog no. 14170112) for 30 min on a rocking platform at RT. The gels were then washed three times with PBS-T for 20 minutes each.

### Pan-ExM sample mounting

After expansion, the gels were incubated overnight in 30% glycerol (Teknova, catalog no. G1797). The gels were next mounted on Poly-L-Lysine-coated glass-bottomed dishes (35 mm; no. 1.5; MatTek). A clean 18-millimeter diameter Poly-L-Lysine-coated coverslip was put on top of the gels after draining excess glycerol using Kimwipes. The samples were then sealed with a two-component silicone glue (Picodent Twinsil, Picodent, Wipperfürth, Germany). After the silicone glue hardened (typically 20-30 min), the samples were stored at dark and at RT until they were imaged.

### Image acquisition

Confocal and STED images were acquired using a Leica SP8 STED 3X equipped with a SuperK Extreme EXW-12 (NKT Photonics) pulsed white light laser as an excitation source and Onefive Katana-08HP pulsed laser as a depletion light source (775-nm wavelength). All images were acquired using either a HC PL APO 63×/1.40-0.60 oil objective or a Plan Apochromat (HC PL APO) 100×/1.40 NA oil immersion CS2 objective. Application Suite X software (LAS X; Leica Microsystems) was used to control imaging parameters. ATTO532 was imaged with 532-nm excitation. ATTO594 was imaged with 585-nm excitation and 775-nm depletion wavelengths.

ATTO647N was imaged with 647-nm excitation and 775-nm depletion wavelengths. DY643 was imaged with 643-nm excitation. SYTOX Green and MitoTracker Orange were excited by 488-nm and 555-nm excitation light, respectively.

Widefield images to measure protein retention were obtained with a Leica tissue culture microscope (DM IL LED FLUO) equipped with a 10×/0.22 NA air objective. An Andor Clara CCD camera operated by MicroManager was used to record images.

### Protein retention assay

HeLa cells were transfected with either GFP-ManII or GFP-Sec61β and plated at 75,000 cells per 12-mm fibronectin-coated glass coverslip. Non-expanded cells from the same experiment were stored in 1× PBS at 4 °C after fixation and were permeabilized with 0.1% Triton X-100 (Sigma-Aldrich, catalog no. T8787) in 1× PBS prior to antibody labeling. All non-expanded, once- and twice-expanded samples were subjected to the same antibody labeling scheme described in the antibody labeling after pan-ExM section, but using a ATTO594-conjugated anti-rabbit antibody (1:250; Sigma-Aldrich, catalog no. 77671) instead of ATTO647N-conjugated antibodies. All samples were imaged in deionized water with a widefield Leica tissue culture microscope (DM IL LED FLUO) using a 10×/0.22 NA air objective (see image acquisition section) and using the same LED light intensity. Total fluorescence signal (TSL) for Golgi samples was quantified by multiplying the total area occupied by stained Golgi stacks with the background-corrected mean fluorescence signal in that area. TSL for ER samples was quantified by measuring the total background-corrected mean fluorescence signal per field of view and dividing it by the number of cells in the field of view. TSL was corrected for the different camera exposure times used. Images were processed using FIJI/ImageJ software.

### Image processing

Images were visualized, smoothed, and contrast-adjusted using FIJI/ImageJ or Imspector software. STED and confocal images were smoothed for display with a 0.7 to 2-pixel sigma Gaussian blur. Minimum and maximum brightness were adjusted linearly for optimal contrast. The TOM20, Mitotracker and confocal ManII data sets were corrected for bleedthrough of the NHS ester channel by subtracting a constant fraction of the latter from the former.

Cristae distance measurements were performed using FIJI. 10-pixel thick line profiles were taken approximately perpendicular to the cristae orientation and peak-to-peak distances were extracted from the profiles. The diameter of ER tubules was determined in FIJI using the Point Tool by manually measuring two positions arranged perpendicular to the orientation of clearly discernible tubules and located at the crests of the signal denoting each side of the tubule. The Euclidean distance between them was used as a measure of the ER tubule diameter. The line profiles shown in Figs. 1 and 2 were extracted from the images using Imspector.

### Expansion factor calculation

Images of HeLa cell nuclei in non-expanded and pan-ExM expanded samples stained with Sytox Green (1:3,000) were acquired with a Leica SP8 STED 3X microscope using a HCX PL Fluotar 10x/0.30 dry objective. Average nuclear cross-sectional area was quantified using FIJI/ImageJ software. To determine the expansion factor, the average nuclear cross-sectional area in pan-ExM samples was divided by the average nuclear cross-sectional area of non-expanded samples. Results are summarized in the **Supplementary Table**.

### Statistics and reproducibility

All experiments, except for the protein retention assay and the data presented in Fig. 2c-f, were carried out three times independently. The n values are stated in figure legends. An unpaired two-tailed t-test was used to analyze the protein retention data presented in Supplementary Fig. 3.

## Supplementary Figures and Videos

**Supplementary Figure 1:**
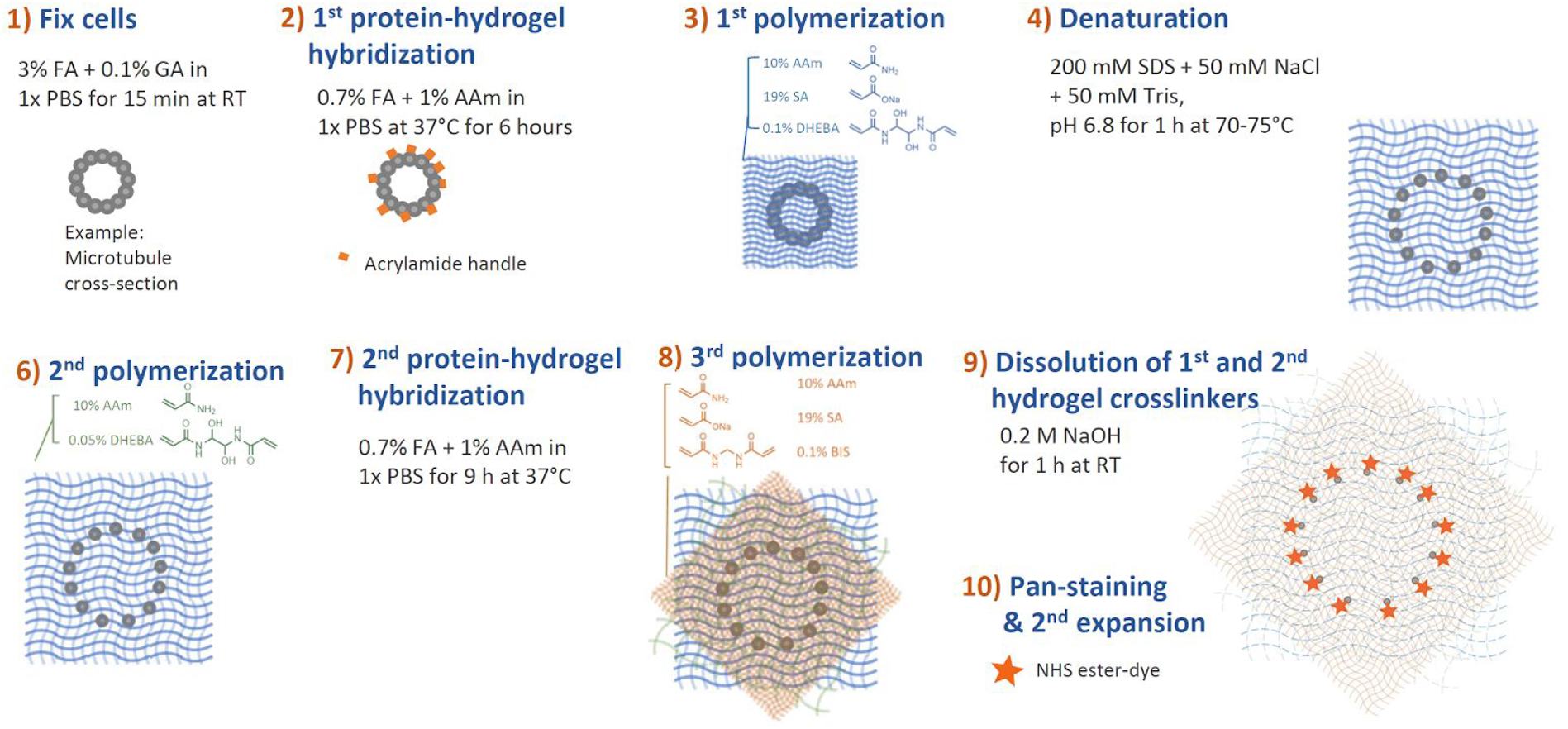
Schematic of pan-ExM

**Supplementary Figure 2:**
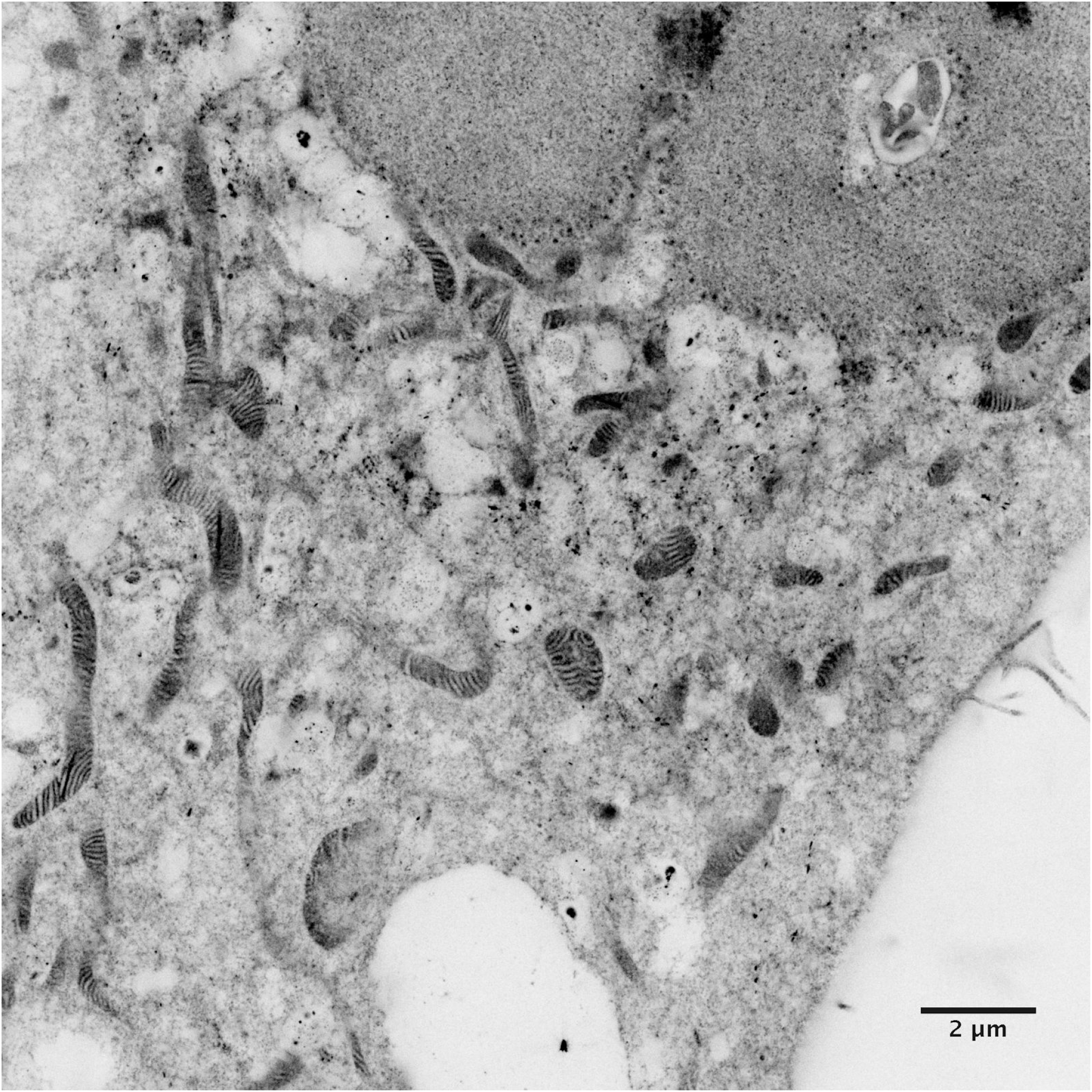
pan-ExM image of a HeLa cell labeled with NHS ester displayed with an inverted color table resembling EM images.

**Supplementary Figure 3:**
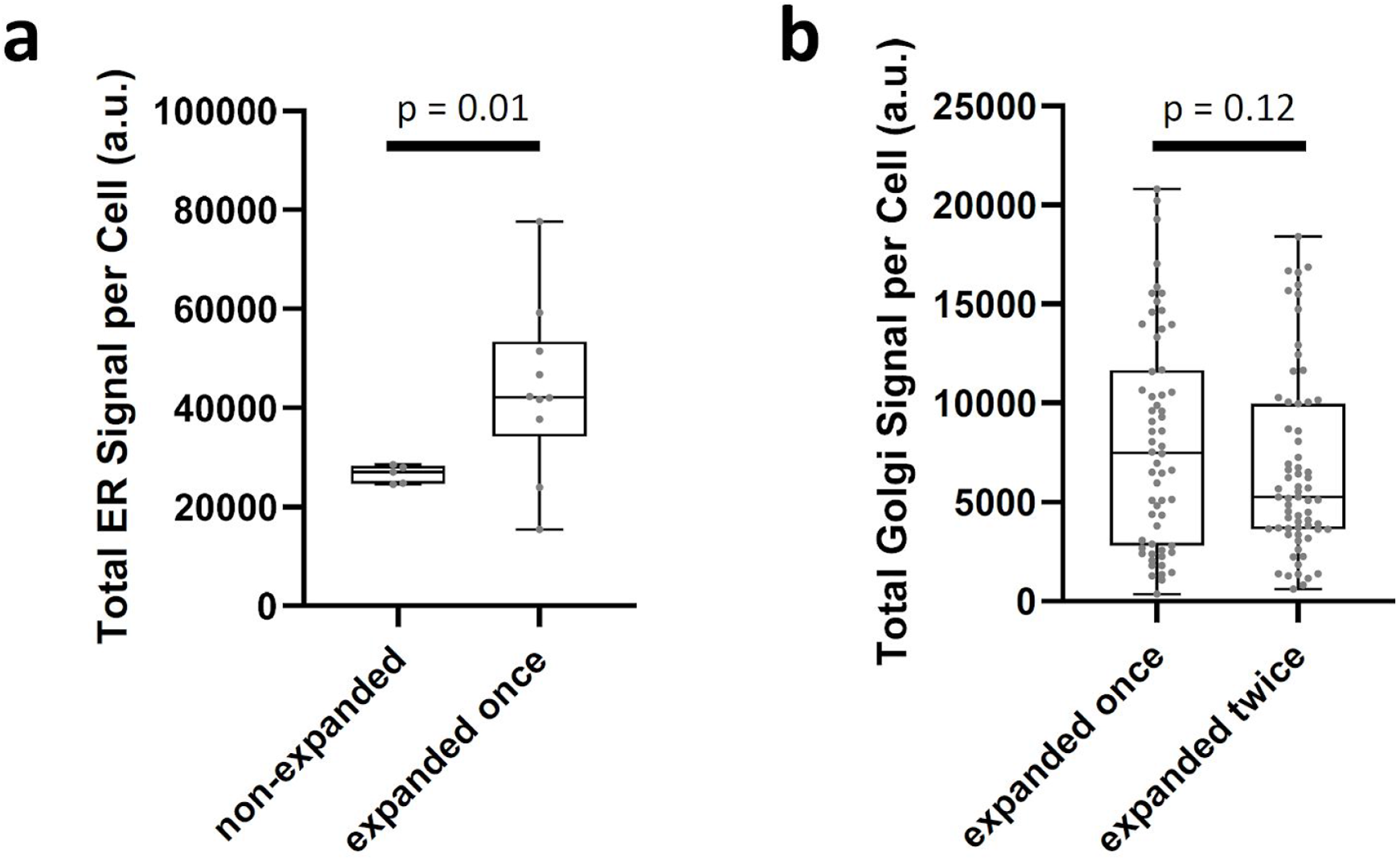
Measurement of protein retention. **a**, comparison of a non-expanded sample with a sample expanded once. *(Non-expanded: n = 2515 cells; Expanded once: n = 294 cells.)* **b**, comparison of a sample expanded once with a sample expanded twice. *(Expanded once: n = 60 cells; Expanded twice: n = 67 cells.) Median and interquartile range are shown with whiskers drawn down to the minimum and maximum values*.

**Supplementary Figure 4:**
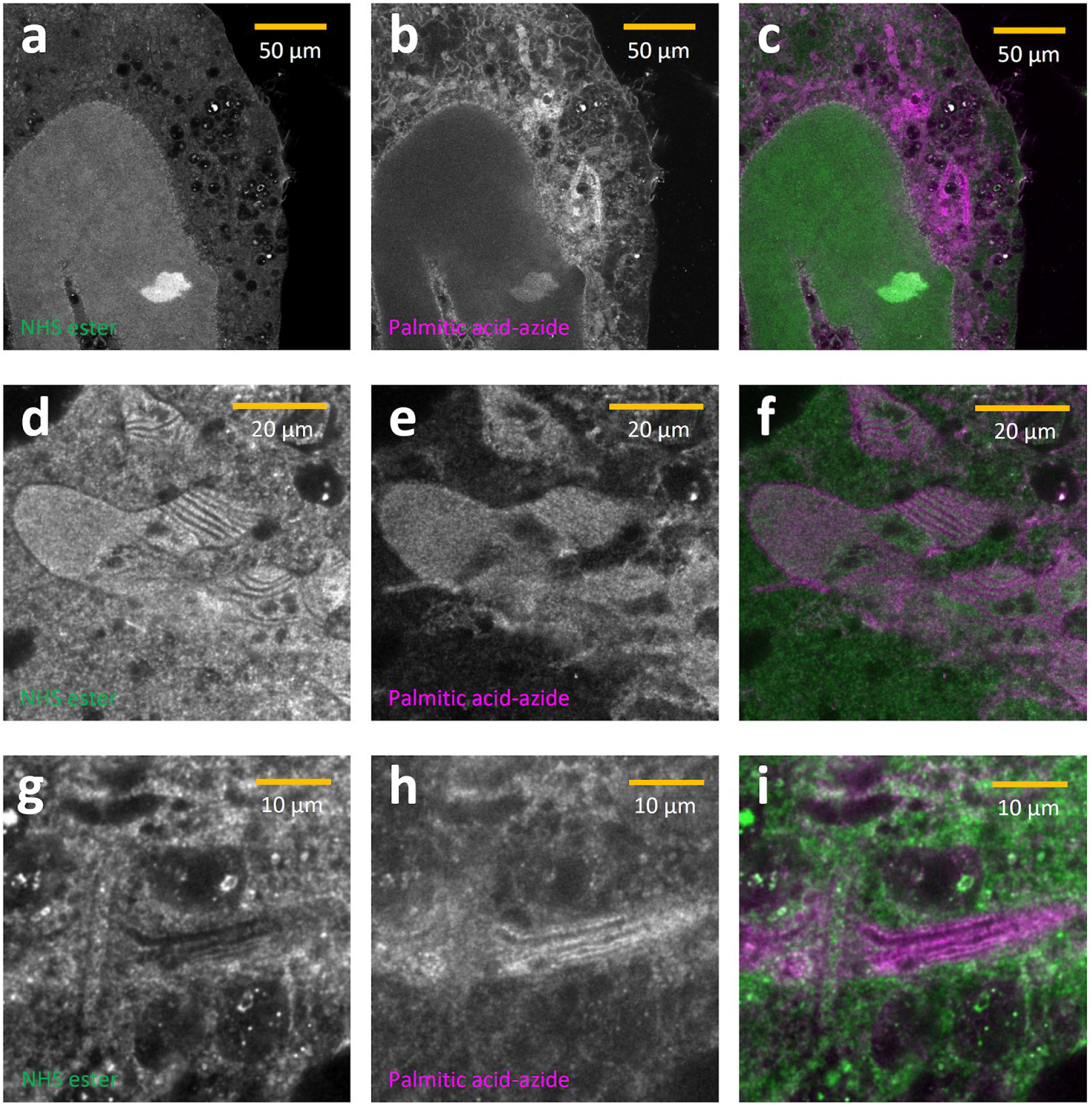
Differential pan-staining of lysine residues and palmitoylated proteins. **a,d,g**, NHS-ester pan-staining highlighting lysine residues. **b,e,h**, Palmitic acid-azide pan-staining highlighting palmitoylated proteins. **c,f,i**, overlay. Scale bars are not corrected for the expansion factor.

**Supplementary Table:** Expansion factors from 5 independent experiments

| **Experiment number** | **Expansion factor** |
| --- | --- |
| Experiment 1 | 14.67 |
| Experiment 2 | 14.33 |
| Experiment 3 | 13.13 |
| Experiment 4 | 13.49 |
| Experiment 5* | 21.03 |

\* The cleavable crosslinker N,N'-Cystaminebisacrylamide (BAC) at a concentration of 0.1% (w/w) was used in the final hydrogel instead of BIS

## Supplementary Video 1

Animation of the data set shown in Figure 2h-s.

## Supplementary Video 2

Animation of a large (186 slices) 3D stack showing a HeLa cell with pan-staining (grey scale) and Mitotracker Orange (fire color table) recorded with a a 63x/1.2 NA water immersion objective on a conventional confocal microscope (Leica TCS SP8).

